# PPAR-γ/PCK1 metabolic pathway modulate synovitis and fibrosis in KOA rats

**DOI:** 10.64898/2026.08.05.742949

**Authors:** Junjie Wu, Xiaodi He, Lucheng Chen, Zhaolin Li, Jie Ling, Hao Xu, Yanwen Hu

## Abstract

**Background:** Knee osteoarthritis (KOA) is a prevalent degenerative joint disease in which synovial inflammation and fibrosis are closely linked to pain, stiffness, and functional limitation. Growing evidence suggests that metabolic dysregulation, particularly in lipid metabolism, is involved in KOA pathogenesis, but the underlying mechanisms remain incompletely defined.

**Methods:** Sprague Dawley rats underwent bilateral anterior cruciate ligament transection to establish a KOA model; sham-operated rats served as controls. RNA sequencing of synovial tissues was performed to identify differentially expressed genes (DEGs) and enriched pathways, followed by GO/KEGG and GSEA analyses. In vivo, adeno-associated virus vectors were used to overexpress or knock down PPAR-γ and phosphoenolpyruvate carboxykinase 1 (PCK1) via intra-articular injection. Ex vivo, primary rat fibroblast-like synoviocytes (FLSs) were stimulated with IL-1β and transfected with PPAR-γ or PCK1 siRNA/overexpression plasmids. synovitis and fibrosis were evaluated by HE, Masson, and Sirius Red staining, immunofluorescence, ELISA, RT-qPCR, and Western blotting.

**Results:** RNA-seq revealed 621 up-regulated and 228 down-regulated genes in KOA synovium versus sham, with DEGs significantly enriched in PPAR signaling, adipocytokine, and AMPK pathways. Metabolism-related genes including *Fabp5, Plin1, Adipoq, Lep, and Pck1* were up-regulated. GSEA indicated downregulation of PPAR-γ signaling in KOA synovium. In vivo and ex vivo, PPAR-γ expression was reduced in KOA, whereas PCK1, FABP5, and ADIPOQ were increased. PPAR-γ overexpression alleviated synovial inflammation, collagen I deposition, and fibrosis, and suppressed FABP5, ADIPOQ, and PCK1 expression; PPAR-γ knockdown produced the opposite effects. Functional studies showed that PCK1 overexpression aggravated synovial inflammatory cell infiltration and fibrosis, elevated IL-1β, IL-18, and TGF-β, and decreased TIMP1 levels in serum, synovial tissue, and FLSs supernatants, whereas PCK1 silencing reversed these changes.

**Conclusions:** The PPAR-γ/PCK1 metabolic axis modulates synovitis and fibrosis in KOA. Downregulation of PPAR-γ and consequent upregulation of PCK1 promote synovitis and fibrotic remodeling. These findings identify the PPAR-γ/PCK1 pathway as a potential therapeutic target for KOA.

## 1. Introduction

Knee osteoarthritis (KOA) is a common degenerative disease with increasing prevalence, primarily attributable to population aging and obesity, and it imposes a substantial burden of disease worldwide ^[1]^. In 2020, more than 100 million people worldwide were estimated to have KOA, most of whom were aged 30 years or older, according to a systematic analysis for the Global Burden of Disease Study ^[2]^. The principal clinical manifestations of KOA include joint swelling and pain, reduced mobility, and, in advanced cases, stiffness and deformity ^[3]^. To date, the pathological mechanisms underlying KOA have yet to be fully elucidated, consequently, effective therapies for mitigating symptoms in established disease remain limited ^[4,5]^.

The major pathological alterations in KOA comprise cartilage loss, subchondral bone structural damage, synovitis, and fibrosis ^[3]^. Earlier studies emphasized cartilage damage as the key driver of KOA. However, in the two decades since the landmark histopathological classification system for synovitis developed by Krenn ^[6]^ et al. in 2002, accumulating evidence suggests that synovial inflammation may be the initiating factor ^[7]^. This chronic, low-grade inflammation, which is strongly associated with innate immunity, contributes to both radiographic progression and pain progression in KOA. It also underlies the excessive production of pro-inflammatory mediators, including members of the interleukin (IL) family and tumor necrosis factor-α (TNF-α), within the synovium and neighboring tissues ^[8]^. In addition, it should be emphasized that when the synovium is immunologically stimulated, fibroblast-like synoviocytes (FLSs) are generally regarded as the primary effector cells, as they constitute the majority of the cellular composition of synovial tissue ^[9]^.

In a healthy joint, the synovium covers the inside of the joint capsule and cavity and has two layers: the intima (the thin lining) and the subintima (the deeper layer). The intima comprises a sparse cellular layer embedded within an amorphous matrix enriched in collagens, notably types I and III, which are the primary constituents responsible for maintaining the extracellular matrix (ECM) of the synovium ^[10]^. Synovitis drives the accumulation of collagen I in the synovium, which is the fundamental cause of synovial fibrosis and is closely associated clinically with joint stiffness and limited function in KOA ^[11]^. Accumulating evidence has demonstrated the critical role of transforming growth factor-β (TGF-β) in fibrotic response of KOA ^[12]^. Besides, in single-cell RNA sequencing (RNA-Seq) of KOA synovial cells, CD34^hi^ fibroblasts were associated with the fibrotic phenotype, and were influenced by mural and endothelial cells ^[13]^. procollagen-lysine, 2-oxoglutarate 5-dioxygenase 2 and collagen type I α1 chain were found to be upregulated, whereas tissue inhibitor of metalloproteinases 1 (TIMP1) was decreased in osteoarthritis-related fibrosis, collectively, these genes are widely considered fibrogenic markers ^[14]^.

On the other hand, given the link between chronic inflammation and metabolic dysregulation, recent studies increasingly highlight the pivotal role of metabolic disturbances in the onset and progression of KOA ^[15,16]^, which may help explain why obesity is a major risk factor for the disease. Machine-learning models identified increased fatty acid intake and reduced saturated fat consumption as strategies to modulate inflammation and improve clinical outcomes in KOA ^[17]^. Furthermore, PPAR-γ signaling, as a central pathway in lipid metabolism, has also been shown to be involved in synovial inflammation in KOA ^[18,19]^, although its downstream effector pathways have not yet been fully elucidated. Meanwhile, Cao ^[20]^ et al. identified an osteoarthritis subtype characterized by disordered synovial lipid metabolism and dysfunction of FLSs, manifested by upregulated expression of perilipin-1 (PLIN1) and fatty acid-binding protein 5 (FABP5). Notably, FABP4, PLIN1 and adiponectin (ADIPOQ) are among the downstream targets through which PPAR-γ mediates inflammatory responses ^[21]^.

While a detailed discussion of the correlation between molecular dysregulation and metabolism in KOA is complex, advances in sequencing technologies have nonetheless provided a means to pursue this line of enquiry. Several metabolism-related genes, including PLIN2 and APOC1, have been identified as differentially expressed genes (DEGs) in KOA ^[22]^. Moreover, the enrichment of these DEGs across distinct signaling pathways provides further insights into the mechanisms of KOA. In this study, we first performed RNA-seq and identified metabolism-related differentially expressed genes in KOA. The upregulation of phosphoenolpyruvate carboxykinase 1 (PCK1) in the synovial tissue of KOA experimental animals may be associated with suppression of PPAR-γ signaling. Subsequently, through in vivo and ex vivo studies, we further demonstrated that the PPAR-γ/PCK1 metabolic pathway modulates synovitis and fibrosis in KOA rats.

## 2. Material and methods

### 2.1 Animals

Male Sprague Dawley rats 180-20 g were purchased from Charles River Laboratories. The animals were maintained in a specific pathogen-free laminar-flow environment and provided with a standard laboratory pellet diet. Housing conditions were carefully controlled, including a temperature of 25 ± 2 °C, relative humidity of 55 ± 5%, and a 12 h light/dark cycle. All animal protocols were reviewed and approved by the Animal Management Committee and the Animal Ethics Committee of Nanjing University of Chinese Medicine (Approval No. 202502A036). All experimental procedures were performed in accordance with the National Institutes of Health Guide for the Care and Use of Laboratory Animals.

### 2.2 KOA modeling

Except for the sham group, KOA was induced in all rats according to a previously reported method ^[23]^ by surgical transection of the anterior cruciate ligament (ACL) in both knees. In the sham group, the skin was incised and the joint cavity was exposed, but the ACL was left intact, after which the incision was closed layer by layer.

### 2.3 In vivo intervention

The KOA model was established by day 14 post-operation, upon which intervention in vivo was started. Transcriptional regulation of PCK1 or PPAR-γ was achieved in vivo, according to the method established by Gallage S, et al ^[24]^. Adeno-associated virus (AVV) containing shRNAs against PPAR-γ (Vector biolabs, USA) or PCK1 (Vector biolabs, USA) were administered to KOA rats by intra-articular injection once weekly for 3 weeks, the corresponding groups were designated as sh-PPAR-γ and sh-PCK1. Specifically, 1×10^10^ AAV particles in a 50 µL volume were injected into the knee joint cavity, as described previously ^[25]^. To overexpress PCK1 or PPAR-γ, AAV vectors overexpressing either PPAR-γ or PCK1 was intra-articular injected, and the corresponding groups were designated as ad-PPAR-γ and ad-PCK1. The Sham were treated with AVV containing scrambled shRNA as negative control for the same periods.

### 2.4 Primary culture of FLSs

In brief, synovial tissues of rats were minced into 2 mm² pieces, digested with 0.1% collagenase type II (Sigma, USA) for 30 min, filtered through a cell strainer, and centrifuged at 1,500 rpm for 4 min. The cell pellet was cultured in DMEM with 10% fetal calf serum (Gibco, USA) and antibiotics (100 U/mL penicillin, 100 μg/mL streptomycin, Invitrogen, USA). FLSs at passages 3-6 were used for experiments.

### 2.5 Ex vivo intervention

Small interfering RNA (siRNA) oligonucleotides against PPAR-γ, PCK1, overexpression plasmid PPAR-γ-pcDNA3.1, PCK1-pcDNA3.1 and nontargeting scrambled siRNA or pcDNA3.1 were purchased from Hanbio (Shanghai, China), Lipofectamine 3000 (Thermo Fisher Scientific, USA) was used to transfect 100 nm siRNA in opti-MEM (Thermo Fisher Scientific, USA) following the manufacturer’s protocol. FLSs were treated with IL-1β (10 ng/mL) for 6 h to mimic the inflammatory environment of KOA, and an equal volume of sterile normal saline was administered as the vehicle control.

### 2.6 RNA-seq

RNA-seq was performed on synovial tissues from KOA and Sham groups. Total RNA was extracted using the EZ-press RNA Purification Kit, with purity/concentration assessed by NanoDrop 2000 and integrity by Agilent 2100 Bioanalyzer. Libraries were constructed using the TruSeq Stranded mRNA LT Sample Prep Kit and sequenced on the Illumina HiSeq 3000. DEGs were identified using limma, edgeR, and DESeq2 (|fold change|≥1.5, *P*<0.05). GO enrichment was performed and visualized with OmicShare, and GSEA was conducted using the GSEA software (Broad Institute, USA).

### 2.7 Immunofluorescence

Specimens were prepared as described previously. Following deparaffinization and rehydration, the sections were soaked in citrate buffer (10 mmol/L citric acid, pH 6.0) to unmask the antigen. Then sections were stained with the primary antibody and the fluorescent secondary antibody (Proteintech, China). Nuclei were labeled with 4, 6-diamidino-2-phenylindole (DAPI, Thermo Fisher Scientific, USA) and images were obtained using a confocal microscope (FluoView FV1000, Olympus, Japan).

### 2.8 Histological analysis

Synovial tissues were fixed in 10% neutral buffered formalin, paraffin-embedded, and sectioned at 5 μm. HE, Masson’s trichrome, and Sirius Red staining were performed according to kit instructions, and sections were imaged on a Leica DMI3000B microscope under bright-field illumination.

### 2.9 Enzyme linked immunosorbent assay (ELISA)

The levels of IL-1β, IL-18, TGF-β and TIMP1 in rat serum or FLS supernatants were measured using commercial ELISA kits (Beyotime Biotechnology, China) according to the manufacturer’s instructions.

### 2.10 Real-time quantitative polymerase chain reaction (RT-qPCR)

Total RNA was isolated from tissue and cell pellets with TRIzol Reagent (Takara Biotechnology, Japan) and reverse transcribed with reverse transcription reagent (Takara Biotechnology, Japan) according to the manufacturer’s protocol. Complementary DNA was used for real-time PCR with SYBR Premix Taq (Vazyme Biotech, China) in a Light Cycler. Relative quantification of gene expression was performed with the comparative threshold method. Changes in mRNA expression levels were calculated after normalization to values for the GAPDH calibrator gene. The primers used are shown in Table.1.

**Table 1.**
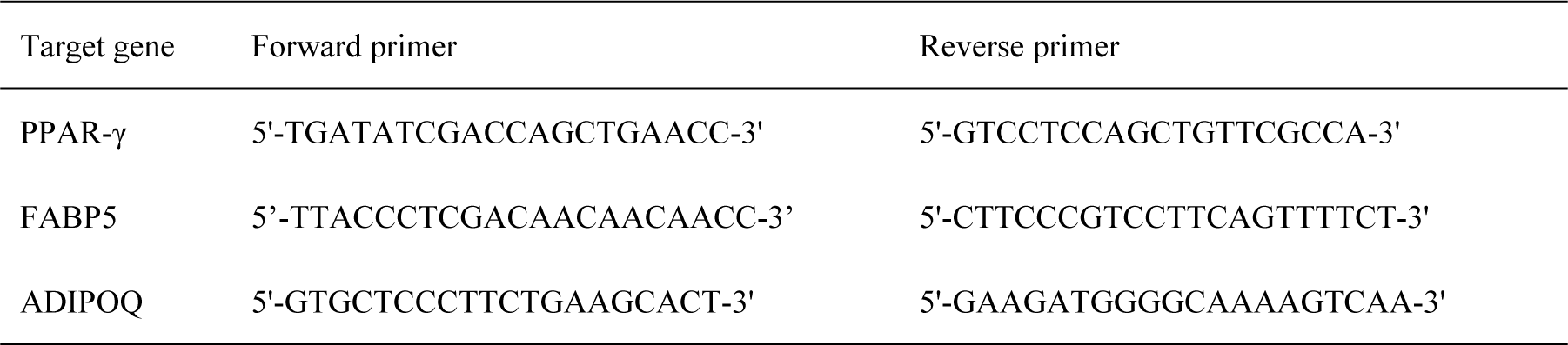
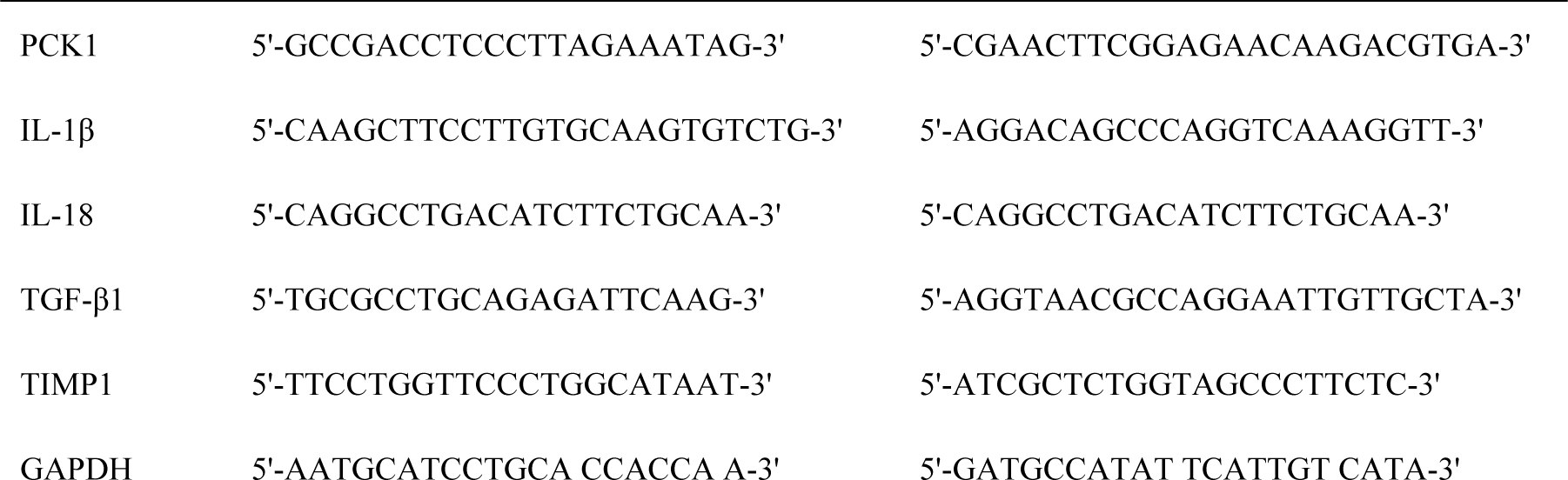
Nucleotide sequences of primers used for qPCR amplification.

### 2.11 Western blotting

Synovial tissues or FLSs were lysed using lysis buffer. The samples were separated by SDS-PAGE for 70 min and the proteins were subsequently transferred to membranes by the wet transfer method. Each membrane was incubated with primary antibodies overnight at 4 °C on a shaker. Following incubation with specific secondary antibodies, the membranes were exposed for visualization.

### 2.12 Statistical analysis

All experiments were performed at least three times. All statistical analyses were performed using SPSS Statistics, Version 28.0 (IBM Corp., USA), and graphs were generated with GraphPad Prism 8.0. For unpaired experiments, two-tailed Student t test, linear regression analysis, or two-way ANOVA was performed. For paired experiments, two-tailed paired t test or linear mixed effect models were utilized. A value of *P*<0.05 was considered as statistically significant.

## 3. Results

### 3.1 RNA-seq reveals downregulation of PPAR-γ signaling in KOA synovial tissue

To better characterize changes in gene expression under inflammatory conditions in KOA synovium, we performed RNA sequencing on synovial tissues from KOA and Sham groups. Principal component analysis (PCA, Fig.1a) showed clear separation of the two groups based on gene expression clustering, and both groups exhibited consistent sequencing depth (Fig.1b). Compared with Sham, KOA group identified 621 up-regulated genes and 228 down-regulated genes. Of note, the metabolism-related genes *Fabp5, Plin1, Adipoq, Lep* and *PCK1* were upregulated (Fig.1c) in the KOA group, in contrast to the Sham. KEGG analysis (Fig.1d, e) indicated that DEGs between the two groups were chiefly enriched in the PPAR signaling pathway, Adipocytokine signaling pathway, and AMPK signaling pathway, a finding corroborated across limma, DESeq2 and edgeR. Furthermore, GSEA plots evaluating genes related to the PPAR (Fig.1f), AMPK (Fig.1g), and TNF signaling pathways (Fig.1h) showed that these gene sets were upregulated in the KOA group and downregulated in the KOA group, except for the TNF signaling pathway, which displayed the opposite trend. Together, RNA-seq revealed downregulation of PPAR-γ signaling in KOA synovial tissue. The DEGs were predominantly metabolism-related genes, and they may be downstream targets of PPAR-γ.

**Figure 1.**
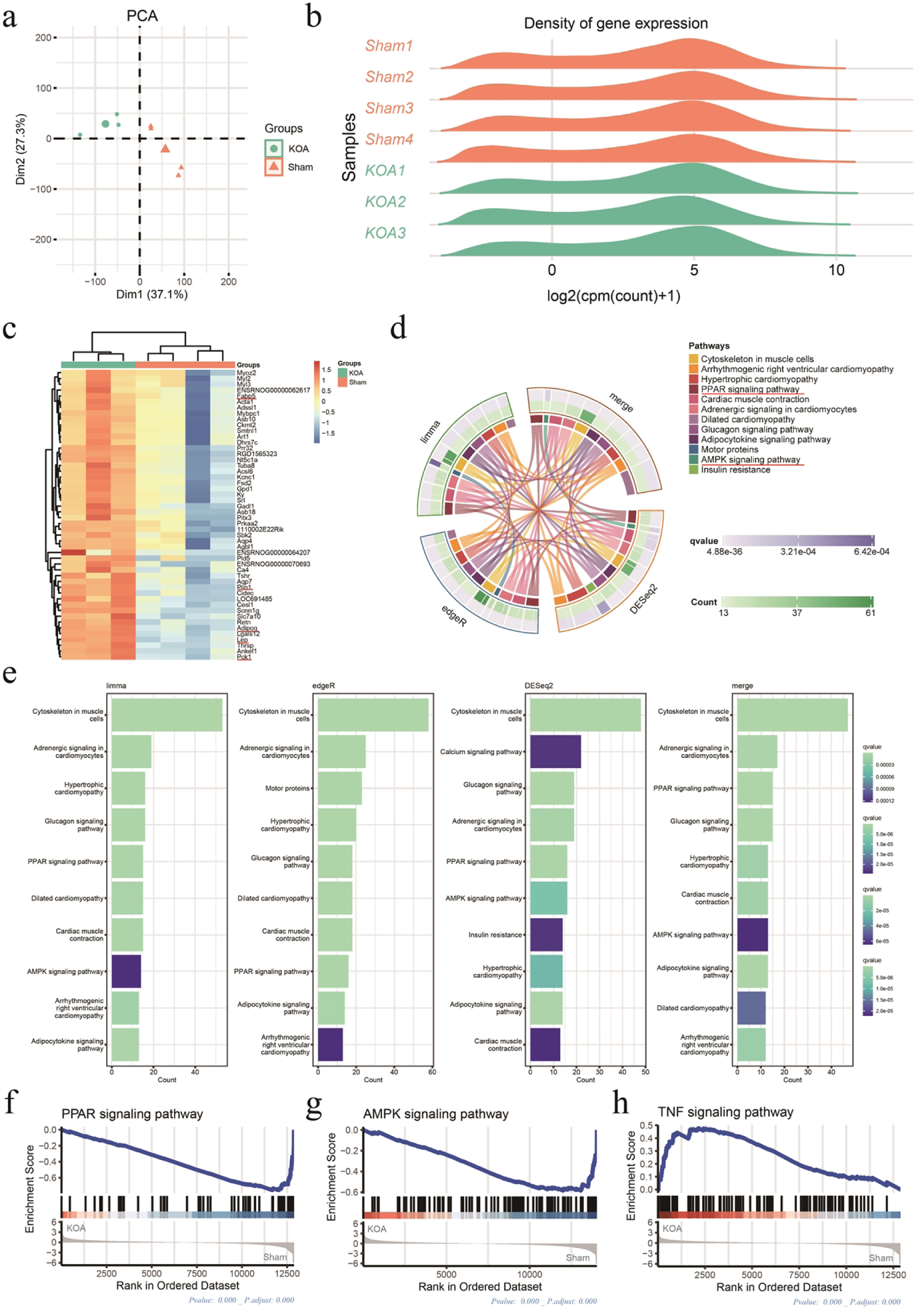
RNA-seq reveals downregulation of PPAR-γ signaling in KOA synovial tissue. Notes: (a). The individual PCA plot of gene expression in synovial tissue from KOA (green) and Sham (orange) groups revealed strong phenotype-specific clustering. (b). Gene expression density for synovial tissues from the two groups of rats. (c). Heatmap of DEGs in synovial tissues from the two groups. (d). String plot revealed KEGG analysis Enrich circle conducted by DEGs between KOA vs. Sham. (e). KEGG analysis of differentially expressed genes identified between the two groups with limma, edgeR, and DESeq2. (f). GSEA analysis of DEGs enriched in PPAR signaling pathway. (g). GSEA analysis of DEGs enriched in AMPK signaling pathway. (h). GSEA analysis of DEGs enriched in TNF signaling pathway.

### 3.2 PPAR-γ signaling modulates metabolism-related genes in synovium in vivo

In the synovial tissue of KOA, larger numbers of inflammatory cells were observed, with disordered cellular arrangement accompanied by more severe synovial fibrosis and type I collagen deposition. Upregulation of PPAR-γ expression by ad-PPAR-γ alleviated synovial inflammation and fibrosis, whereas sh-PPAR-γ silenced synovial PPAR-γ expression and exacerbated the severity of synovitis and fibrosis in KOA (Fig.2a). We further assessed PPAR-γ expression in rat synovial tissue from each group by measuring fluorescence intensity. The fluorescence intensity of PPAR-γ was significantly reduced in the KOA group compared with the Sham group. Intervention with ad-PPAR-γ reversed the KOA-induced decrease in PPAR-γ fluorescence intensity, whereas sh-PPAR-γ further reduced PPAR-γ fluorescence intensity in the KOA group (Fig.2b, c). Subsequently, we examined the relative gene expression of PPAR-γ, FABP5, ADIPOQ and PCK1 in synovial tissues from each group. These genes were enriched in the PPAR-γ signaling pathway and are closely related to metabolism, according to the RNA-seq results. Consistent with the RNA-seq data, the relative gene expression of PPAR-γ was significantly reduced in the KOA group compared with the Sham group, whereas it was significantly increased in the ad-PPAR-γ group compared with the KOA group. In contrast, sh-PPAR-γ further decreased the relative gene expression of PPAR-γ in the KOA group (Fig.2d). Notably, intervention with ad-PPAR-γ significantly suppressed the KOA-induced increase in FABP5, ADIPOQ and PCK1 gene expression in synovial tissue, whereas sh-PPAR-γ led to a markedly greater upregulation of FABP5, ADIPOQ and PCK1 expression compared with the KOA group (Fig.2e-g). As expected, the relative protein expression levels of PPAR-γ, FABP5, ADIPOQ and PCK1 in synovial tissues from each group showed the same pattern of differences as their relative gene expression levels(Fig.2h-l). In summary, these data indicate that PPAR-γ signaling modulates the expression of metabolism-related genes in the synovium in vivo.

**Figure 2.**
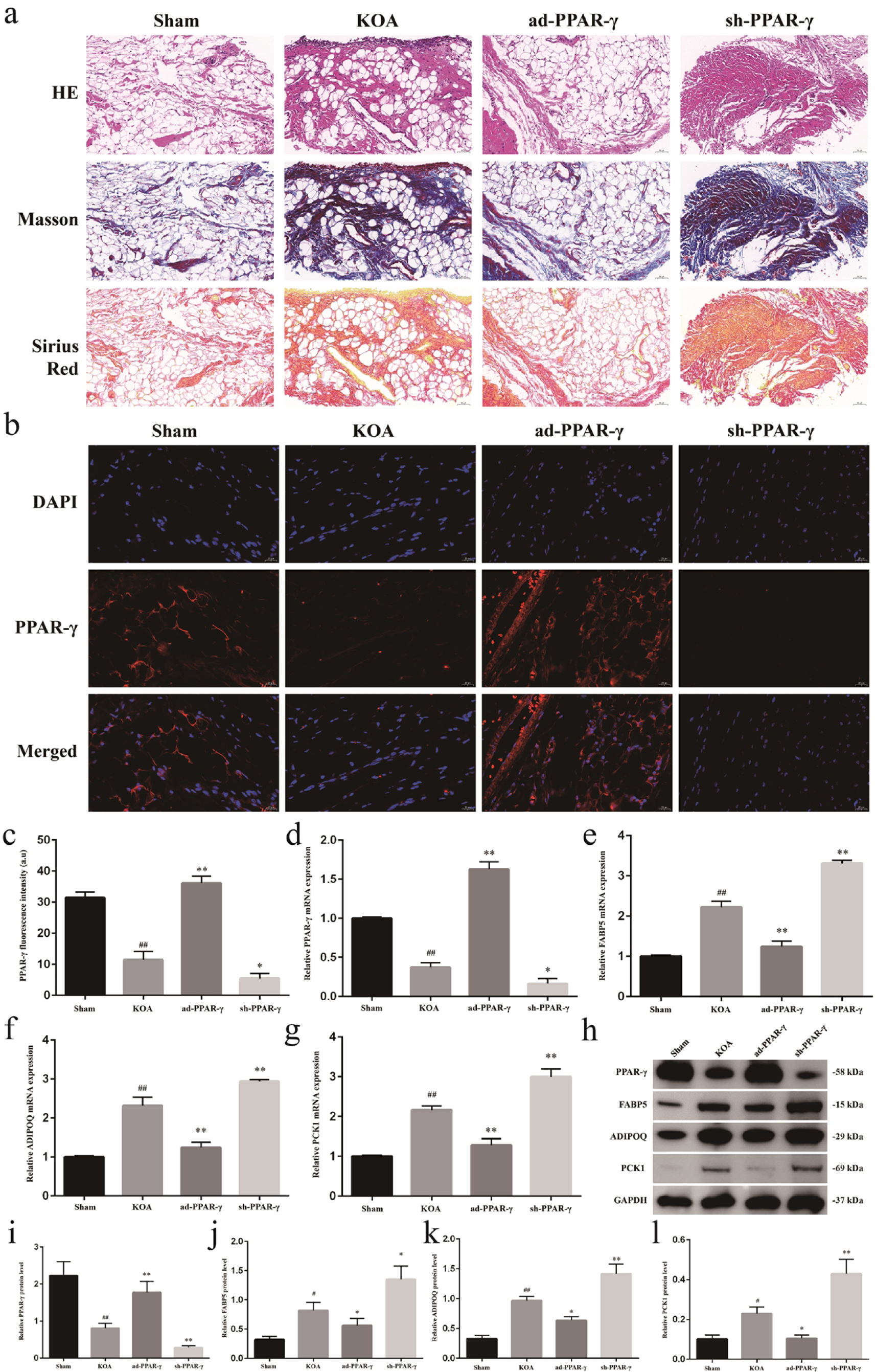
PPAR-γ signaling modulates metabolism-related genes in synovium in vivo. Notes: (a). Representative HE, Masson’s trichrome, and Sirius Red stained sections of synovial tissue from each rat group. 200x, scale bar=50μm. (b). Representative immunofluorescence staining of PPAR-γ in synovial tissue. 400x, scale bar=20μm. (c). Quantification of relative PPAR-γ fluorescence intensity in synovial tissue. ^##^*P*<0.01 vs. Sham, \**P*<0.05, \*\**P*<0.01 vs. KOA. (d). Relative mRNA expression of PPAR-γ in synovial tissue. ^##^*P*<0.01 vs. Sham, \**P*<0.05, \*\**P*<0.01 vs. KOA. (e). Relative mRNA expression of FABP5 in synovial tissue. ^##^*P*<0.01 vs. Sham, \*\**P*<0.01 vs. KOA. (f). Relative mRNA expression of ADIPOQ in synovial tissue. ^##^*P*<0.01 vs. Sham, \*\**P*<0.01 vs. KOA. (g). Relative mRNA expression of PCK1 in synovial tissue. ^##^*P*<0.01 vs. Sham, \*\**P*<0.01 vs. KOA. (h). Representative protein bands of PPAR-γ, FABP5, ADIPOQ and PCK1. (i). Protein expression levels of PPAR-γ in synovial tissue. ^##^*P*<0.01 vs. Sham, \*\**P*<0.01 vs. KOA. (j). Protein expression levels of FABP5 in synovial tissue. ^#^*P*<0.05 vs. Sham, \*\**P*<0.05 vs. KOA. (k). Protein expression levels of ADIPOQ in synovial tissue. ^##^*P*<0.01 vs. Sham, \**P*<0.05, \*\**P*<0.01 vs. KOA. (l). Protein expression levels of PCK1 in synovial tissue. ^#^*P*<0.05 vs. Sham, \**P*<0.05, \*\**P*<0.01 vs. KOA.

### 3.3 PPAR-γ signaling modulates metabolism-related genes in FLSs ex vivo

In the ex vivo FLSs model, the fluorescent intensity of PPAR-γ was markedly reduced in the KOA group compared with the Sham group. Transfection with pc-PPAR-γ induced transcriptional upregulation of PPAR-γ in FLSs, resulting in increased fluorescence intensity relative to the KOA group, whereas si-PPAR-γ silenced PPAR-γ expression in FLSs and accordingly reduced its fluorescence intensity (Fig.3a, b). Consistent results were obtained by qPCR, which showed a similar trend in the relative mRNA expression level of PPAR-γ in FLSs (Fig.3c). Furthermore, the expression of the metabolism-related genes FABP5, ADIPOQ and PCK1 was upregulated in the KOA group compared with the Sham. Overexpression of PPAR-γ reversed this KOA-induced upregulation, whereas silencing PPAR-γ led to an even more pronounced increase in their transcript levels relative to the KOA group (Fig.3d-f). This finding is consistent with the above in vivo results, and the protein expression levels of PPAR-γ, FABP5, ADIPOQ and PCK1 in FLSs across the different groups were also in line with their corresponding gene expression levels (Fig.3g-k).

**Figure 3.**
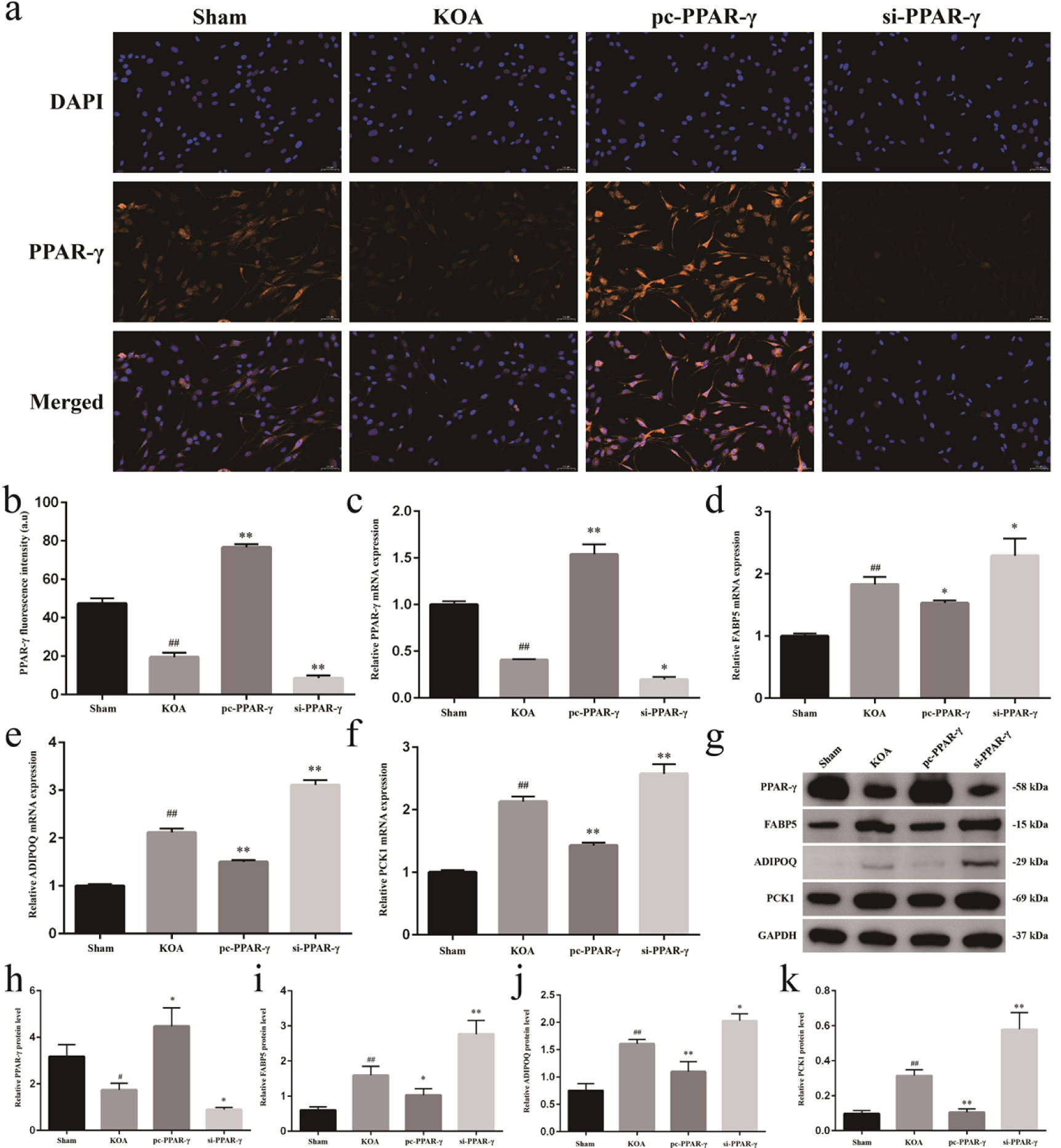
PPAR-γ signaling modulates metabolism-related genes in FLSs ex vivo. Notes: (a). Representative immunofluorescence staining of PPAR-γ in FLSs. 400x, scale bar=20μm. Quantification of relative PPAR-γ fluorescence intensity FLSs. ^##^*P*<0.01 vs. Sham, \*\**P*<0.01 vs. KOA. (c). Relative mRNA expression of PPAR-γ in FLSs. ^##^*P*<0.01 vs. Sham, \**P*<0.05, \*\**P*<0.01 vs. KOA. (d). Relative mRNA expression of FABP5 in FLSs. ^##^*P*<0.01 vs. Sham, \**P*<0.05 vs. KOA. (e). Relative mRNA expression of ADIPOQ in FLSs. ^##^*P*<0.01 vs. Sham, \*\**P*<0.01 vs. KOA. (f). Relative mRNA expression of PCK1 in FLSs. ^##^*P*<0.01 vs. Sham, \*\**P*<0.01 vs. KOA. (g). Representative protein bands of PPAR-γ, FABP5, ADIPOQ and PCK1. (h). Protein expression levels of PPAR-γ in FLSs. ^#^*P*<0.05 vs. Sham, \**P*<0.05 vs. KOA. (i). Protein expression levels of FABP5 in FLSs. ^##^*P*<0.01 vs. Sham, \**P*<0.05, \*\**P*<0.01 vs. KOA. (j). Protein expression levels of ADIPOQ in FLSs. ^##^*P*<0.01 vs. Sham, \**P*<0.05, \*\**P*<0.01 vs. KOA. (k). Protein expression levels of PCK1 in FLSs. ^##^*P*<0.01 vs. Sham, \*\**P*<0.01 vs. KOA.

### 3.4 PPAR-γ attenuates synovial inflammation and fibrosis in KOA via PCK1 in vivo

After elucidating the regulatory role of PPAR-γ signaling on PCK1 in the KOA model, we further sought to determine whether PCK1 influences synovial inflammation and fibrosis during KOA progression. Similarly, we used an adenoviral vector to induce either overexpression or silencing of PCK1 in the synovial tissue of KOA. Histopathological examination of the synovium showed that, compared with the KOA group, the ad-PCK1 group exhibited more pronounced inflammatory cell infiltration, disordered cellular arrangement, and more severe collagen deposition and synovial fibrosis, whereas sh-PCK1 alleviated synovial inflammation and delayed the progression of fibrosis in KOA (Fig.4a). Immunofluorescence results not only revealed the upregulation of PCK1 in the synovial tissue of KOA, but also confirmed the in vivo efficacy of the adenovirus, as PCK1 exhibited higher relative fluorescence intensity in the ad-PCK1 group than in the KOA group, whereas its fluorescence intensity was markedly reduced in the sh-PCK1 group compared with the KOA group (Fig.4b, c).

**Figure 4.**
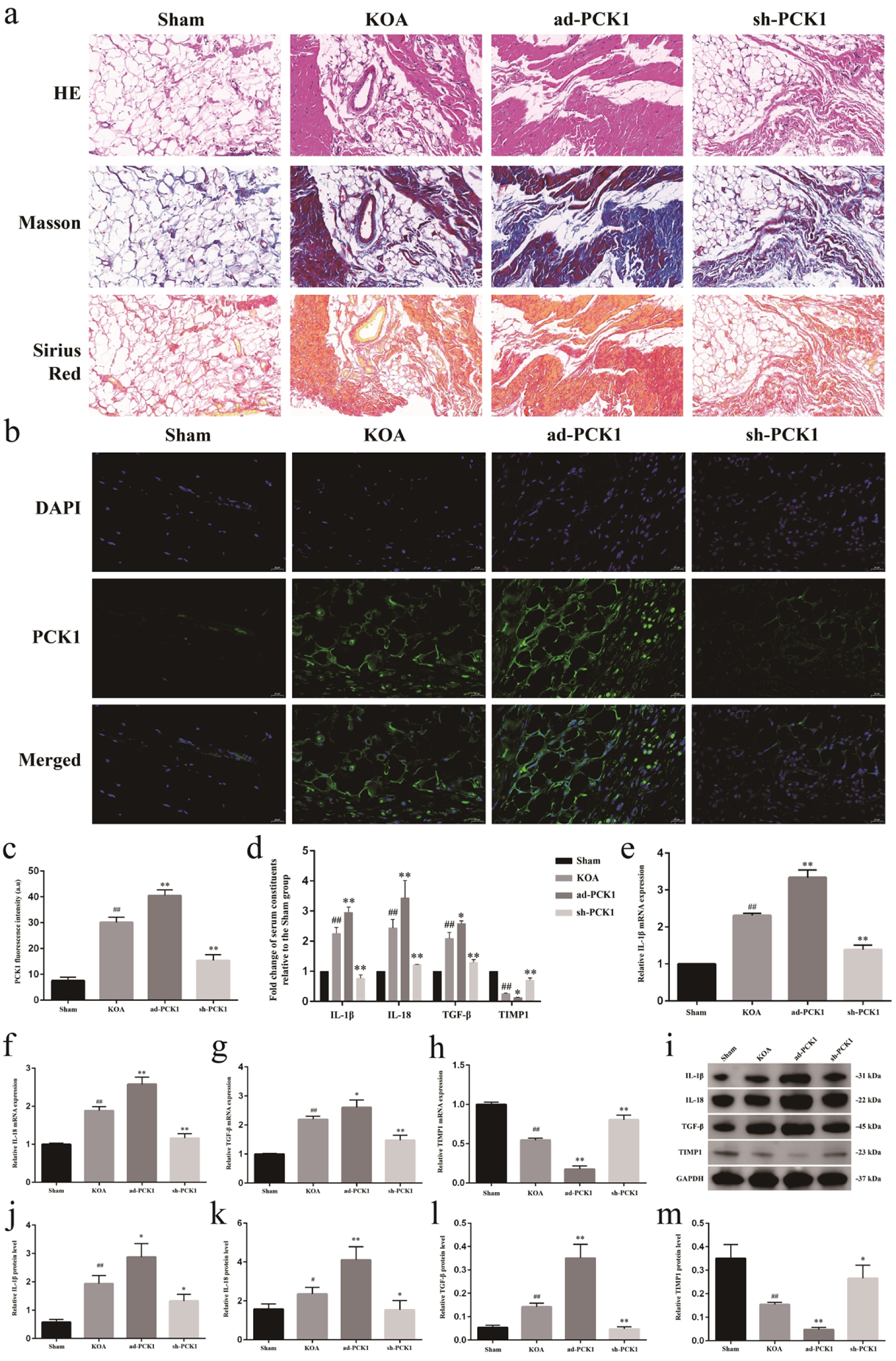
PPAR-γ attenuates synovial inflammation and fibrosis in KOA via PCK1 in vivo. Notes: (a). Representative HE, Masson’s trichrome, and Sirius Red stained sections of synovial tissue from each rat group. 200x, scale bar=50μm. (b). Representative immunofluorescence staining of PCK1 in synovial tissue. 400x, scale bar=20μm. (c). Quantification of relative PCK1 fluorescence intensity in synovial tissue. ^##^*P*<0.01 vs. Sham, \*\**P*<0.01 vs. KOA. (d). The serum levels of IL-1β, IL-18, TGF-β and TIMP1 in rats relative to those in Sham. ^##^*P*<0.01 vs. Sham, \**P*<0.05, \*\**P*<0.01 vs. KOA. (e). Relative mRNA expression of IL-1β in synovial tissue. ^##^*P*<0.01 vs. Sham, \*\**P*<0.01 vs. KOA. (f). Relative mRNA expression of IL-18 in synovial tissue. ^##^*P*<0.01 vs. Sham, \*\**P*<0.01 vs. KOA. (g). Relative mRNA expression of TGF-β in synovial tissue. ^##^*P*<0.01 vs. Sham, \**P*<0.05, \*\**P*<0.01 vs. KOA. (h). Relative mRNA expression of TIMP1 in synovial tissue. ^##^*P*<0.01 vs. Sham, \*\**P*<0.01 vs. KOA. (i). Representative protein bands of IL-1β, IL-18, TGF-β and TIMP1. (j). Protein expression levels of IL-1β in synovial tissue. ^##^*P*<0.01 vs. Sham, \**P*<0.05 vs. KOA. (k). Protein expression levels of IL-18 in synovial tissue. ^##^*P*<0.01 vs. Sham, \**P*<0.05, \*\**P*<0.01 vs. KOA. (l). Protein expression levels of TGF-β in synovial tissue. ^##^*P*<0.01 vs. Sham, \*\**P*<0.01 vs. KOA. (m). Protein expression levels of TIMP1 in synovial tissue. ^#^*P*<0.05 vs. Sham, \**P*<0.05, \*\**P*<0.01 vs. KOA.

Furthermore, we measured the serum levels of IL-1β, IL-18, TGF-β and TIMP1 in each group of rats. Compared with the Sham group, the KOA group showed markedly increased levels of IL-1β, IL-18 and TGF-β, and a reduced level of TIMP1, indicating more severe inflammation and fibrosis in KOA rats. Overexpression of PCK1 further increased the levels of IL-1β, IL-18 and TGF-β and decreased TIMP1, whereas knockdown of PCK1 reversed this trend (Fig.4d). We also examined the mRNA (Fig.4e-h) and protein (Fig.4i-m) expression levels of the above pro-inflammatory cytokines and fibrotic markers in rat synovial tissues, and found that their trends were highly consistent with the intergroup differences observed in serum levels.

### 3.5 PPAR-γ attenuates synovial inflammation and fibrosis in KOA via PCK1 ex vivo

Finally, we examined whether PCK1 affects the expression of pro-inflammatory cytokines and fibrotic markers in the ex vivo KOA model. The immunofluorescence intensity of PCK1 (Fig.5a, b) was significantly increased in FLSs from the KOA model, and intervention with pc-PCK1 and si-PCK1 further enhanced and reduced PCK1 fluorescence intensity, respectively. The contents of IL-1β, IL-18 and TGF-β in the FLS supernatant were markedly higher in the KOA group than in the Sham group, whereas TIMP1 levels were reduced. pc-PCK1 further increased the supernatant levels of IL-1β, IL-18 and TGF-β and decreased TIMP1 in KOA FLSs, while si-PCK1 reduced IL-1β, IL-18 and TGF-β levels and increased TIMP1 levels (Fig.5c). Meanwhile, the intergroup differences in the relative mRNA (Fig.5d-g) and protein (Fig.4h-l) expression of the above pro-inflammatory cytokines and fibrotic markers in FLSs were highly consistent with the intergroup differences observed in their levels in the FLSs’ supernatant. Taken together with the upstream regulatory effect of PPAR-γ signaling on PCK1 expression described above, these results validate that PPAR-γ attenuates synovial inflammation and fibrosis in KOA via PCK1.

**Figure 5.**
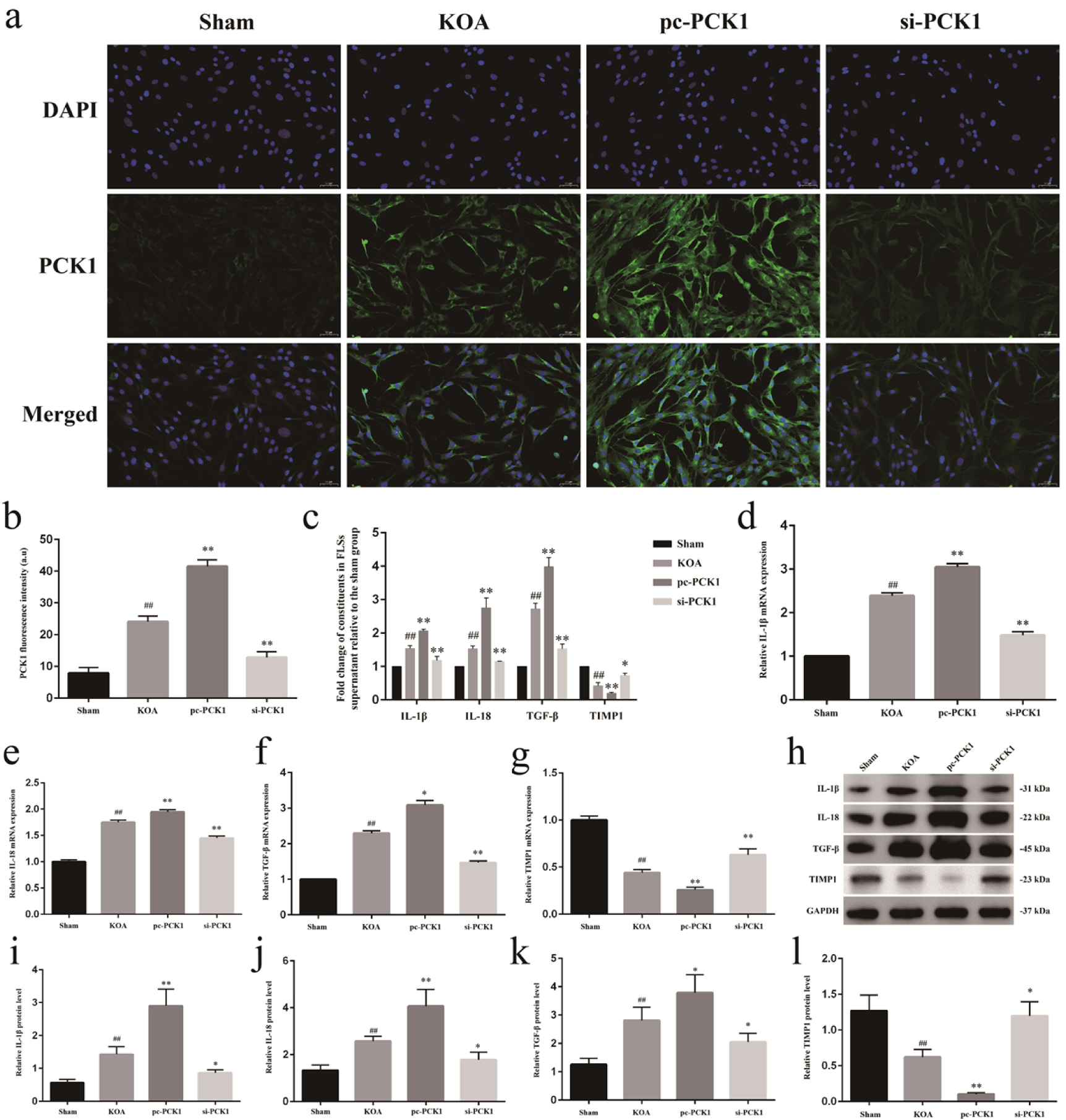
PPAR-γ attenuates synovial inflammation and fibrosis in KOA via PCK1 ex vivo. Notes: (a). Representative immunofluorescence staining of PCK1 in FLSs. 400x, scale bar=20μm. (b). Quantification of relative PCK1 fluorescence intensity FLSs. ^##^*P*<0.01 vs. Sham, \*\**P*<0.01 vs. KOA. (c). The levels of IL-1β, IL-18, TGF-β and TIMP1 in the supernatant of FLSs relative to the Sham group. ^##^*P*<0.01 vs. Sham, \**P*<0.05, \*\**P*<0.01 vs. KOA. (d). Relative mRNA expression of IL-1β in FLSs. ^##^*P*<0.01 vs. Sham, \*\**P*<0.01 vs. KOA. (e). Relative mRNA expression of IL-18 in FLSs. ^##^*P*<0.01 vs. Sham, \*\**P*<0.01 vs. KOA. (f). Relative mRNA expression of TGF-β in FLSs. ^##^*P*<0.01 vs. Sham, \**P*<0.05, \*\**P*<0.01 vs. KOA. (g). Relative mRNA expression of TIMP1 in FLSs. ^##^*P*<0.01 vs. Sham, \*\**P*<0.01 vs. KOA. (h). Representative protein bands of IL-1β, IL-18, TGF-β and TIMP1. (i). Protein expression levels of IL-1β in FLSs. ^##^*P*<0.01 vs. Sham, \**P*<0.05, \*\**P*<0.01 vs. KOA. (j). Protein expression levels of IL-18 in FLSs. ^##^*P*<0.01 vs. Sham, \**P*<0.05, \*\**P*<0.01 vs. KOA. (k). Protein expression levels of TGF-β in FLSs. ^##^*P*<0.01 vs. Sham, \**P*<0.05 vs. KOA. (l). Protein expression levels of TIMP1 in FLSs. ^##^*P*<0.01 vs. Sham, \**P*<0.05, \*\**P*<0.01 vs. KOA.

## 4. Discussion

Metabolic factors in the onset and progression of KOA have attracted increasing attention in recent years, and the metabolic phenotype is even regarded as an independent subtype of KOA. Here, the first pathway to be highlighted is the AMPK signaling pathway ^[26]^. As a common upstream regulator of multiple metabolic activities, including energy metabolism, AMPK has been shown to be closely associated with inflammation and senescence phenotypes in KOA ^[27,28]^, as well as with the balance between cartilage synthesis and degradation ^[29]^. The classical AMPK agonist metformin has already been shown, in clinical studies and high-quality basic research, to exert a beneficial effect on the progression of KOA ^[30,31]^. PPAR-γ, meanwhile, is an important downstream effector of AMPK in the regulation of lipid metabolism ^[32]^.

PPARs belong to the nuclear receptor family and are classified as intracellular receptors that function as transcription factors upon activation ^[33]^. To date, three types of PPARs, namely PPAR-α, PPAR-δ and PPAR-γ, have been identified, and PPAR-γ ligands activate insulin-sensitizing genes that are mainly involved in adipogenesis, metabolism and inflammation ^[33]^. Besides, PPAR-γ senses incoming non-esterified long-chain fatty acids (LCFAs) and induces pathways that store LCFAs as triglycerides.

In this process, ADIPOQ, may act an important PPAR-γ target to maintain their metabolic activity and insulin sensitivity ^[21]^. FABP5 is another prototypical gene induced by PPAR-γ. It is abundantly expressed in adipocytes and binds to LCFAs with high affinity ^[34]^, and it plays an important role in lipid metabolism in KOA. Xu et al. demonstrated that inhibiting CC chemokine receptor 1 reduced the inflammatory response, alleviated cartilage ageing and slowed degeneration via the PPAR-γ signaling pathway. Chen et al. used RNA-seq to identify that tenascin-N protein is up-regulated in KOA, and that it accelerates inflammation and cartilage damage in KOA by impairing AMPK/PPAR-γ signaling ^[35]^. In this study, we first performed RNA-seq on synovial tissues from KOA model animals to identify metabolism-related DEGs and enriched pathways. Consistent with the above findings, we observed that the metabolism-related genes *Fabp5, Plin1, Adipoq, Lep*, and *PCK1* were upregulated in the KOA group and were predominantly enriched in the PPAR-γ signaling pathway.

We also verified the regulatory role of PPAR-γ signaling in synovial lipid metabolism in KOA through both in vivo and ex vivo experiments. Adenoviral overexpression or knockdown of PPAR-γ respectively suppressed or enhanced the mRNA and protein expression levels of the metabolism-related genes FABP5, ADIPOQ and PCK1. Among these genes, PCK1 particularly attracted our attention. PCK1 is the first rate-limiting enzyme in gluconeogenesis and converts oxaloacetate to phosphoenolpyruvate in the cytoplasm ^[36]^. PPAR-α and PCK1 may act cooperatively in the process of hepatic fibrogenesis ^[37]^. Furthermore, the upregulation of PCK1 in tissues may promote the expression of the pro-inflammatory cytokine TNF-α, thereby exacerbating inflammation ^[38,39]^. In this study, we first reported the upregulated expression of PCK1 in the synovial tissue of KOA rats. Overexpression of PCK1 further increased the levels of the pro-inflammatory cytokine IL-1β and the pro-fibrotic factor TGF-β, while suppressing the expression of the anti-fibrotic factor TIMP1 in KOA synovium. In contrast, silencing PCK1 exerted beneficial anti-inflammatory and anti-fibrotic effects in KOA rats.

Synovial inflammation and fibrosis have attracted increasing attention in the field of KOA, as they are closely associated with the clinical symptoms of pain and joint stiffness ^[40]^. Among these, TGF-β, as a central pro-fibrotic marker, plays a double-edged role ^[41]^. It must be noted that TGF-β, as an essential factor for growth and development, also exerts beneficial effects in KOA treatment, including tissue repair, maintaining bone homeostasis and preserving the stem cell niche ^[42,43]^. However, its pro-fibrotic and pro-angiogenic actions, which disrupt the synovial ECM microenvironment, should likewise be emphasized.

However, our study still has several limitations. First, due to experimental constraints, the number of sequencing samples was limited. In future work, increasing the sample size or performing joint analyses with similar sequencing datasets will undoubtedly enhance the robustness of our conclusions. Second, our current investigation of PCK1 is restricted to the level of expression. Subsequent studies will focus on its enzymatic activity as a metabolic enzyme, aiming to elucidate in greater depth the specific metabolic alterations through which it regulates inflammation and fibrosis in KOA.

### Conclusions

In summary, this study demonstrates that PPAR-γ induced metabolic alterations modulate synovitis and fibrosis in KOA rats through PCK1. Upregulation of PCK1 accelerates synovial inflammation and fibrotic progression in KOA. These results provide a mechanistic foundation for the PPAR-γ/PCK1 metabolic pathway and highlight a promising therapeutic target for KOA management. Future studies should explore its long-term efficacy in human trials and delineate its interplay with specific lipid metabolic pathways.

## Ethics approval and consent to participate

The animal study protocol was approved by the Animal Management Committee and Animal Ethics Committee of Nanjing University of Chinese (Approval No. 202502A036), and all experiments were conducted in strict accordance with the National Institutes of Health Guidelines for the Care and Use of Laboratory Animals.

## Authors’ contributions

JJW: methodology, validation, investigation, resources, writing—original draft preparation; XDH: methodology, supervision; LCC: formal analysis, investigation, validation; ZLL: formal analysis, investigation; JL: investigation; HX: writing—review and editing, supervision, project administration; YWH: validation data curation, project administration, funding acquisition.

## Acknowledgements

The authors would like to thank the Nanjing Vane Biotechnology Co., for their help.

## Availability of data and material

The data used to support the findings of this study are available from the corresponding author upon request.

